# Integrative assessment of total and intact HIV-1 reservoir by a five-region multiplexed Rainbow digital PCR assay

**DOI:** 10.1101/2023.08.18.553846

**Authors:** Mareva Delporte, Willem van Snippenberg, Evy E. Blomme, Sofie Rutsaert, Maxime Verschoore, Evelien De Smet, Marie-Angélique De Scheerder, Sarah Gerlo, Linos Vandekerckhove, Wim Trypsteen

## Abstract

Persistent latent reservoirs of intact HIV-1 proviruses, capable of rebounding despite suppressive ART, hinder efforts towards an HIV-1 cure. Hence, assays specifically quantifying intact proviruses are crucial to assess the impact of curative interventions. Clinical trials have utilized two recent assays: intact proviral DNA assay (IPDA) and Q4PCR. While IPDA is more sensitive due to amplifying short fragments, it may overestimate intact fractions by relying only on two small regions. Q4PCR is sequencing-based and its performance might be subjected to bias against full-length proviruses. Leveraging digital PCR (dPCR) advancements, we developed the ‘Rainbow’ 5-plex proviral HIV-1 DNA assay, assessing it with standard materials and samples from 69 PLWH. The Rainbow assay proved equally sensitive but more specific than IPDA, is not subjected to bias against full-length proviruses, enabling high-throughput quantification of total and intact reservoir size. This innovation offers potential for targeted evaluation and monitoring of rebound-competent reservoirs, contributing to HIV-1 management and cure strategies.

**Teaser:** The 5 color ‘HIV-1 Rainbow’ digital PCR assay offers a multi-level view on the HIV reservoir in one snapshot reaction.

## Introduction

The human immunodeficiency virus (HIV) remains a major global public health issue with an estimated 38.4 million people living with HIV-1 (PLWH) in 2021 [1]. Although current antiretroviral therapy (ART) regimens can successfully block HIV-1 replication and transmission, the development of a broad and definite HIV-1 cure has proven to be a major challenge due to persisting latent reservoirs. These latent HIV-1 reservoirs contain both intact and defective proviruses, of which the intact, replication-competent proviruses will inevitably reinitiate viral replication after ART is discontinued [2]. In addition, while defective proviruses cannot cause viral rebound, they have the potential to elicit immune activation which causes ongoing inflammation in PLWH [3].

Over the past decades, several HIV-1 clinical trials have been evaluating new therapeutic interventions that hopefully can eliminate or allow for a functional control of the HIV-1 reservoir in PLWH. To monitor the efficacy of these curative efforts, it is crucial that HIV-1 cure studies implement a precise and accurate method to quantify the HIV-1 reservoir [4, 5]. Importantly, the clinically relevant reservoir, composed of intact, replication-competent proviruses constitutes only a fraction of the total reservoir. To specifically quantify this intact reservoir, either cell culture-based techniques (VOA, TILDA, HIV-Flow, VIP-spot) or PCR-based techniques are used, that generally lead to respectively an under- or overestimation of the intact reservoir. The cell culture-based methods, are often labor-intensive and fail to reflect the true viral reservoir size due to the inability of reactivating all latent HIV-1 viruses [6–10]. PCR-based assays, including the total HIV-1 DNA assay, intact proviral DNA assay (IPDA), 5T-IPDA and cross-subtype IPDA (CS-IPDA), rely on the amplification of multiple target regions in the HIV-1 genome to quantify HIV-1 DNA while Q4PCR requires an additional sequencing step to quantify HIV-1 DNA [4, 5, 11–13].

The total HIV-1 DNA assay is widely used in HIV-1 clinical trials to monitor total HIV-1 DNA levels as biomarker for the size of the viral reservoir but is unable to distinguish between intact and defective proviruses [10, 11]. The IPDA is now considered the go-to assay to quantify the intact viral reservoir in ART-treated PLWH. Although proven as a powerful approach, the IPDA still overestimates the intact viral reservoir, as this assay is not able to detect all large deletions between psi and *env* [12]. This overestimation, due to the misclassification of non-intact sequences as intact, complicates the evaluation of the effect of curative interventions [2, 14–16]. Therefore, the estimation/quantification of intact proviruses can be improved by implementing more target regions to better annotate the correct intact sequences [16].

The Q4PCR was a first attempt to increase the number of included intactness regions and integrates a quantitative PCR approach using four probes with near full-length genome sequencing (nFGS) for sequence verification of intact and defective genomes, resulting in an estimate of the fraction of intact proviruses [5]. Even though the Q4PCR is a valuable method, it is less high-throughput than IPDA due to the requirement for a prior limiting-dilution *gag* qPCR and full-length genome amplification before an intactness readout is performed. Moreover, recent studies show that nFGS methods fail to amplify the majority of full-length proviruses, inducing quantification bias. This could lead to a significant underestimation of the frequency of intact proviruses, making it difficult to assess the effect of curative interventions [2, 10, 17].

Moreover, a clinically relevant assay is necessary that offers sufficient throughput, demands a relatively small amount of biological sample and maximizes information retrieval on the (intact) viral reservoir. In this light, we took advantage of the current evolution in digital PCR (dPCR) platforms that offer an increased multiplex capacity and designed the “Rainbow proviral HIV-1 DNA dPCR assay” to simultaneously gather information on total and intact proviral HIV-1 DNA levels and provide a more accurate intactness result. This Rainbow HIV-1 DNA assay targets five HIV-1 subgenomic regions by combining three existing PCR-based assays in one multiplex dPCR set-up: the total HIV-1 DNA assay, the intact proviral DNA assay (IPDA) and the quadruplex qPCR (Q4PCR) assay [5, 11, 12]. Hence, the Rainbow HIV-1 DNA assay serves as an intermediary, bridging the gap between IPDA and Q4PCR as this Rainbow assay is more specific than IPDA and is not subjected to bias against full-length proviruses as in Q4PCR.

In this study, the Rainbow proviral HIV-1 DNA dPCR assay was optimized on reference standard material and performed on a cohort of 69 PLWH (subtype B). In conclusion, the Rainbow proviral HIV-1 DNA dPCR assay demonstrated comparable sensitivity while exhibiting enhanced specificity compared to IPDA and is not subjected to bias against full-length proviruses. The Rainbow assay can serve as a high-throughput assay that is successful in capturing different levels of information on the viral reservoir size in a single reaction, offering potential for targeted evaluation and monitoring of rebound-competent reservoirs, contributing to HIV-1 management and cure strategies.

## Results

### Designing the five-region Rainbow proviral HIV-1 DNA dPCR assay

The Rainbow proviral HIV-1 DNA dPCR assay is designed to capture five regions of the HIV-1 genome simultaneously by combining a readout for total HIV-1 DNA and intact HIV-1 DNA (*Figure 1*). The selected intactness regions, which are located in relatively conserved parts of the HIV-1 genome, including the packaging signal (psi), envelope (*env*), group-specific antigen (*gag*) and polymerase (*pol*), were previously identified through in-silico analysis of full-length clade-B sequences and were selected for their ability to discriminate between intact and defective proviruses. Furthermore, all target regions are based on published primers and probes [5, 12, 18, 19]. These primers and probes were adapted to work in a 5-plex dPCR reaction on the QIAcuity Four platform by testing out different combinations of primer/probe concentrations and fluorophores on J-Lat 10.6 gDNA (cells that are latently infected with a proviral clone of HIV-1 subtype B). In the end, one combination was selected that resulted in accurate quantification and an optimal resolution to facilitate thresholding (*Table S1, Figure S1-S2).* Signal spillover from adjacent channels was deduced for each fluorescence channel by performing a monocolor control experiment where only one fluorescent channel at a time has positive partitions. Here, signal spillover was observed in the orange channel, coming from the red channel (*Figure S3*). This spillover was taken into account when setting the thresholds manually in the Qiacuity Software Suite.

**Figure 1:**
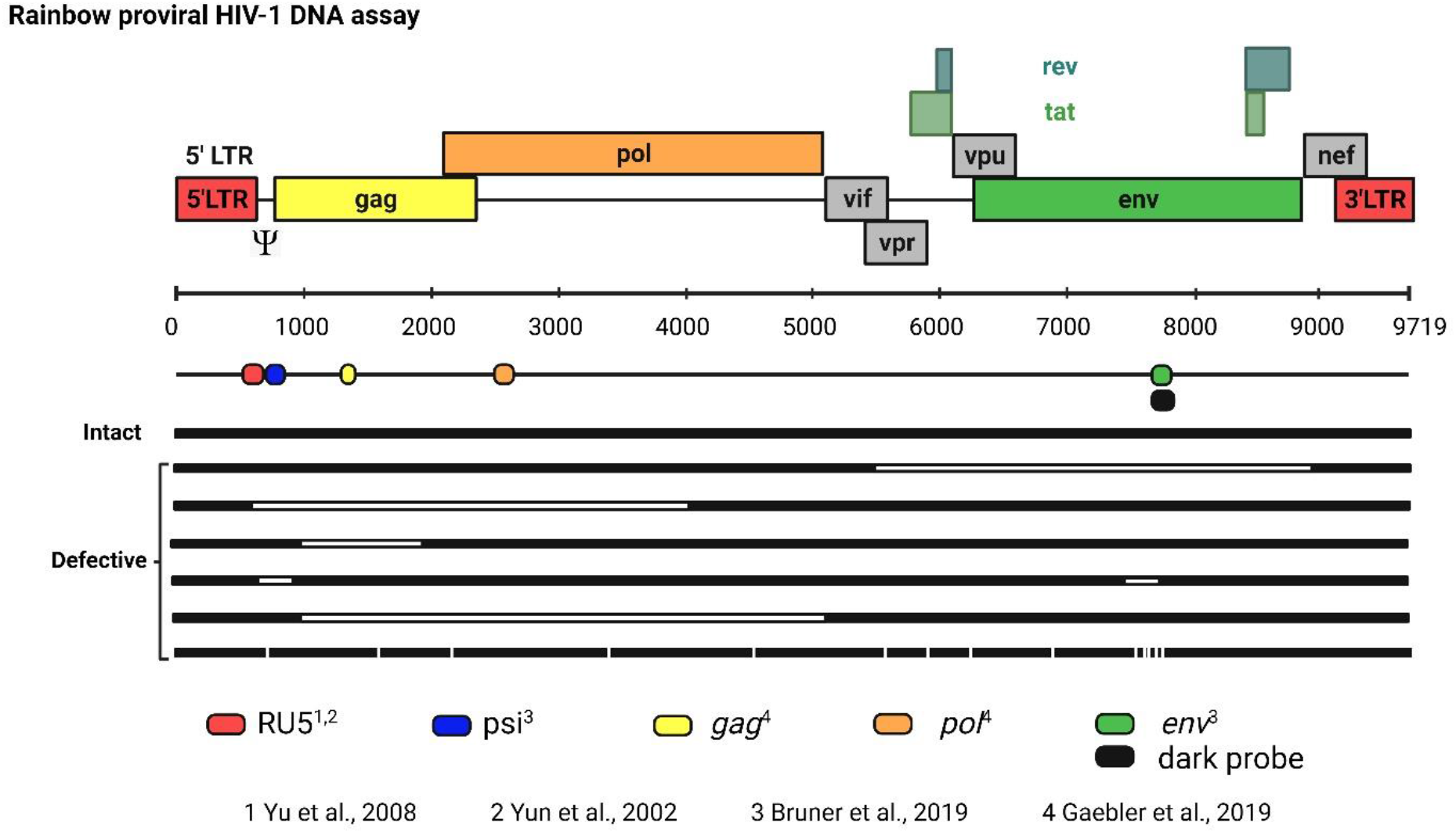
Schematic overview of the Rainbow proviral HIV-1 DNA dPCR assay. The five target regions used in this assay are displayed at their respective locations in the HIV-1 genome, with a different color for each region. RU5 is used to quantify total HIV-1 DNA, whereas the other four regions (psi, *gag*, *pol* and *env*) are used to quantify intact HIV-1 DNA. A dark probe without fluorescent dye is used at the same location of *env* to bind hypermutated *env* (competition probe). (created with www.Biorender.com)

Next, the technical performance of the Rainbow HIV-1 DNA assay was assessed in-depth on three independent standard curves that encompass a two-fold dilution of 12 points with each dilution point measured in 20 replicates. Specificity was tested in 288 NTC samples, containing gDNA from HIV-1 negative CD4+ T cells.

In this study, total HIV-1 DNA is defined by the RU5 target region. Intactness levels were defined by three different combinations of HIV-1 subgenomic regions from the Rainbow HIV-1 DNA assay. When intact HIV-1 DNA was quantified based on two target regions (psi and *env*), the intact copy number is called ‘IPDA positive’. The two other combinations for intact HIV-1 DNA comprise four target regions (psi, *env*, *gag* and *pol*) or five target regions (psi, *env*, *gag*, *pol* and RU5) and are called ‘D4PCR positive’ or ‘Rainbow positive’, respectively.

### Performance of the Rainbow proviral HIV-1 DNA dPCR assay on a standard dilution curve

#### Linearity and precision

Three independent standard dilution curves were prepared containing J-Lat gDNA performance of the Rainbow HIV-1 DNA assay, mimicking samples from PLWH. In terms of quantification, the Rainbow assay resulted in comparable copy numbers for all five multiplexed regions in all dilution points. Linear quantification was observed (R² > 0.99) for all individual regions and for intact HIV-1 DNA, which was quantified based on two, four and five regions (IPDA positive, D4PCR positive and Rainbow positive) and corrected for DNA shearing (*Figure S4)*. The variability between replicates (n = 60) was in the same range for each dilution point per standard dilution curve with a CV% below 20% in the 5 highest dilution points in each standard dilution curve. The CV% increased towards the lower dilution points, as expected *(Figure 2, Table S2*).

**Figure 2:**
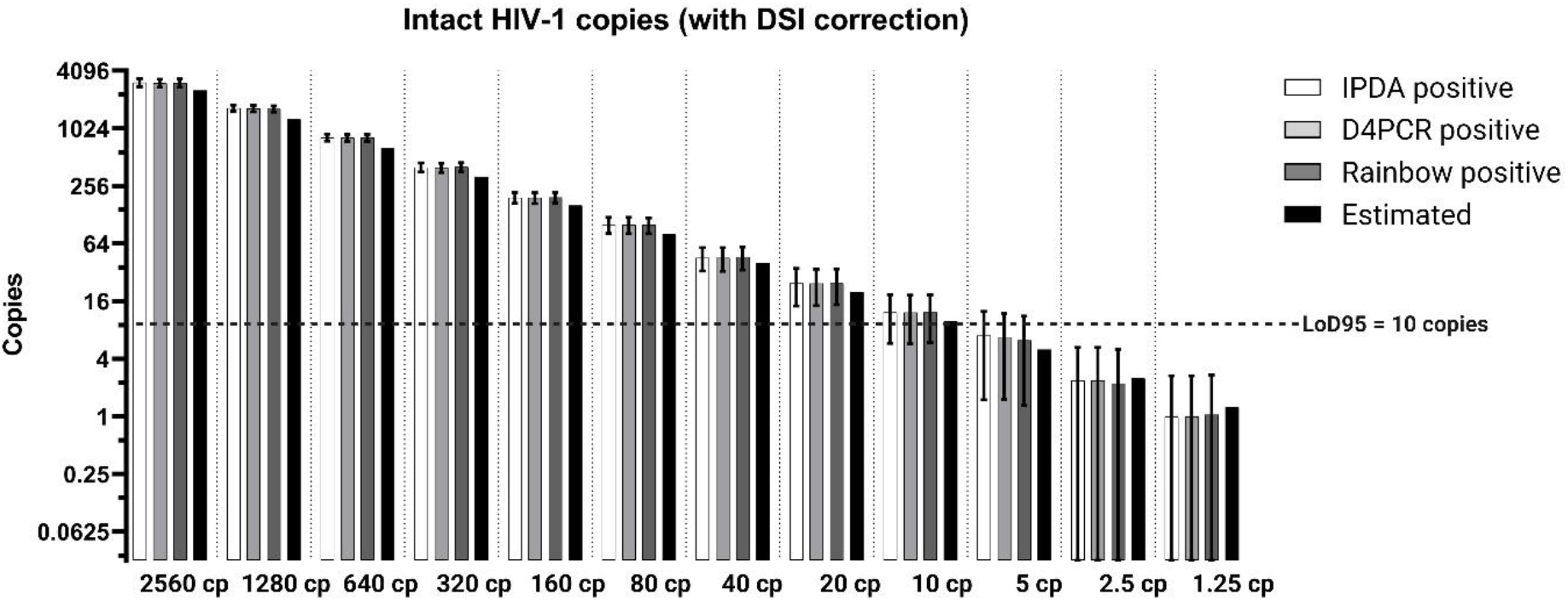
Technical performance of the Rainbow proviral HIV-1 DNA dPCR assay in standard material. Observed intact HIV-1 copies defined by two (IPDA positive), four (D4PCR positive) or five regions (Rainbow positive) for each dilution point (n = 60) and corrected for shearing; Limit of Detection (LoD95) was observed at 10 copies input in the dPCR reaction, corresponding to 0.4 cp/µL stock solution.

#### Sensitivity and specificity

The sensitivity (Limit of Detection) was defined as the lowest DNA input where HIV-1 DNA could be reliably detected in 95% of the replicates (LoD95), as is mentioned in ISO20395 [20]. The LoD95 for each specific target region and for intact HIV-1 DNA (IPDA positive, D4PCR positive and Rainbow positive) was determined at 10 copies gDNA input per reaction as we were able to detect 10 HIV-1 copies in 57/60 replicates (*Figure S5*). The specificity or limit of blank (LoB) of the Rainbow dPCR assay was tested in 288 NTCs of HIV-1 negative CD4 T cell gDNA (∼500 ng per replicate). Only 2 out of 288 replicates showed 1 double-positive partition for psi and *env* (IPDA positive) and none of these replicates were positive for intact HIV-1 DNA defined by four (D4PCR positive) or five (Rainbow positive) regions, resulting in LoB values of 0.32, 0 and 0 intact copies respectively. The LoB for the target regions RU5, psi*, env, gag* and *pol* was 1.27, 1.32, 2.76, 1.30 and 1.32 copies respectively.

### Performance of the Rainbow proviral HIV-1 DNA dPCR assay in PLWH

#### Cohort characteristics

The study cohort included 69 individuals infected with HIV-1 subtype B. All included PLWH have an undetectable viral load (<50 copies/mL) and have been receiving suppressive ART for 8 years on average. Other clinical and virological parameters are summarized in *Table 1*.

**Table 1:** Descriptive statistics for the study population of 69 ART-surpressed PLWH. Median frequencies (with interquartile ranges) are shown below, except for age (absolute frequencies with percentage).

| Descriptive Characteristic | Total<br>(N = 69) |
| --- | --- |
| Male (%)* | 65 (94.2%) |
| Age (years) | 48.6 (15) |
| Nadir CD4+ T (cells/μL) | 324 (238) |
| CD4 count (cells/μL) | 700 (303) (N = 68) |
| CD4/CD8 ratio | 1 (0.6) (N = 68) |
| Time from start ART to first time undetectable viremia (months) | 4 (3) (N = 65) |
| ART initiation (months post infection) | 4 (15) (N = 63) |
| Time on ART (years) | 8.3 (5.5) (N = 67) |

#### Rainbow proviral HIV-1 DNA dPCR assay performance

After extensively testing the Rainbow proviral HIV-1 DNA dPCR assay on standard material, the Rainbow HIV-1 DNA assay was used to quantify total and intact HIV-1 DNA in 69 PLWH. Overall, HIV-1 DNA was detected in all 69 samples from PLWH. However, loss of signal (no positive partitions above the LoB) was observed in one or multiple HIV-1 regions in some PLWH. For the individual assays targeting *env*, psi, *gag*, *pol* and RU5, there is detection in 78% (54/69), 90% (62/69), 90% (62/69), 91% (63/69) and 100% of the participants respectively (*Figure 3a*).

**Figure 3:**
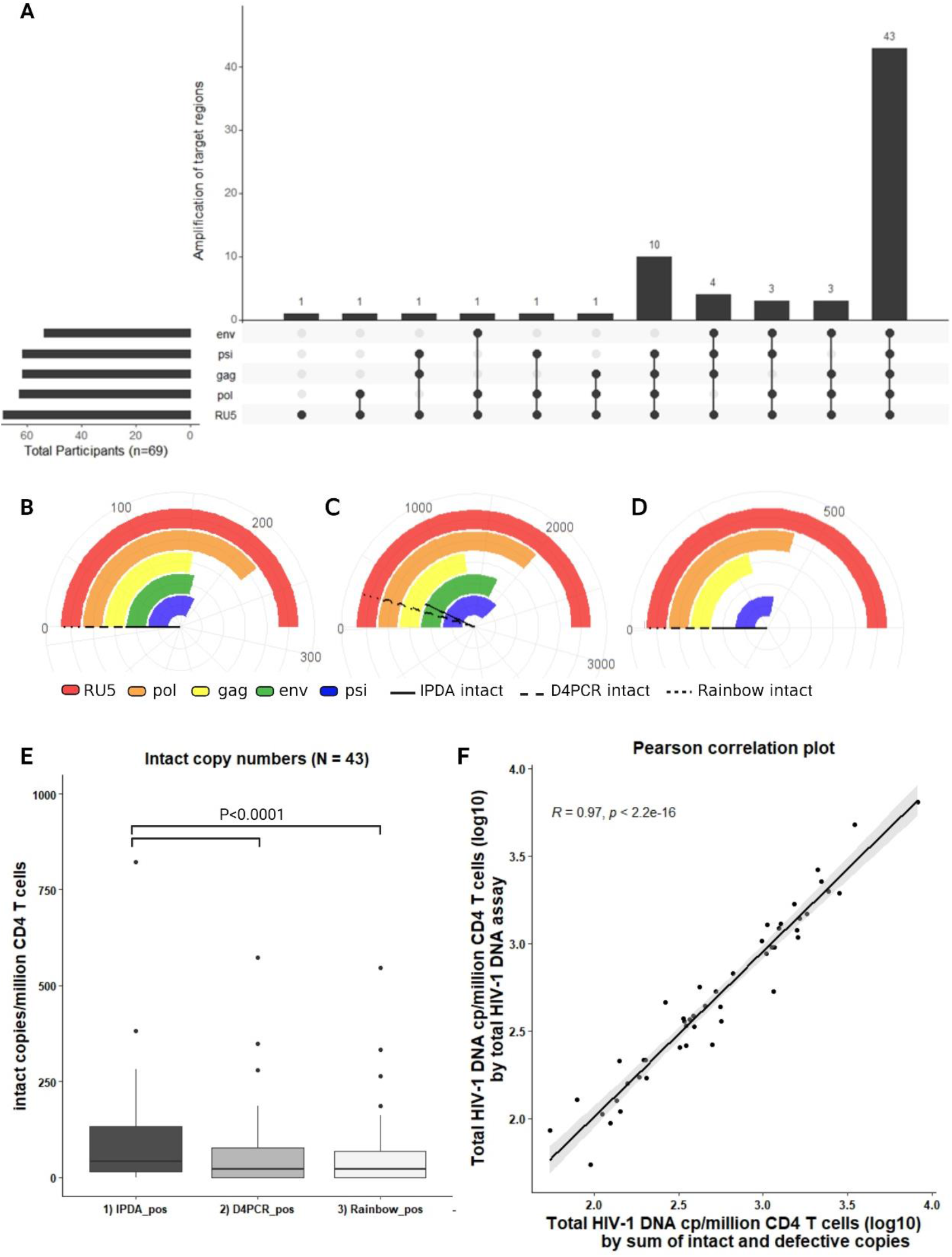
Performance of the Rainbow proviral HIV-1 DNA assay in samples from PLWH. (A) An Upset plot with the amplification results of five target regions of the Rainbow proviral HIV-1 DNA assay in PLWH (n = 69) [31, 32]. The horizontal bars depict in how many participants each target region was successful. The vertical bars demonstrate the number of participants where a combination of different target regions was successful. A grey dot means that there was a loss of signal for that region, a black dot means that partitions were picked up above the limit of blank. (B-D) Rainbow plots (circularized bar charts) of three different PLWH; each color represents the copies/million CD4+ T cells of that target region, the (dashed) lines represent the intactness levels (copies/million CD4+ T cells) quantified by two (IPDA intact), four (D4PCR intact) or five (Rainbow intact) regions. (B) Rainbow plot of a PLWH in whom all target regions were detected but no intactness results were obtained (undetectable intact HIV-1 DNA). (C) Rainbow plot of a PLWH in whom all target regions were detected and intactness results were obtained. (D) Rainbow plot of a PLWH in whom a loss of signal was observed in the env region, leading to an unreportable intactness result. (E) Boxplot of intact copy numbers defined by two, four or five regions in samples from PLWH where no loss of signal occurred (N = 43). The Wilcoxon-signed rank test was used to test the statistical significance of the difference between intactness results obtained by two regions (psi/env) or four/five regions (psi/env/gag/pol/RU5). (F) Pearson correlation plot of total HIV-1 DNA results calculated by the sum of intact and defective copies (IPDA) vs. total HIV-1 DNA results quantified by the total HIV-1 DNA assay.

In a scenario where there are positive partitions for all individual assays, indicating positive PCR reactions, but no double, quadruple or quintuple partitions, we conclude an undetectable finding for intactness (Figure 3b). Quantifiable intactness results are obtained when at least one double, quadruple or quintuple partition is detected (Figure 3c). If a signal failure was observed for a certain assay while the other assays show positive partitions, we define this as an unreportable finding for a certain intactness level readout (*Figure 3d*).

#### Intact HIV-1 DNA

Similar to the quantification of intact HIV-1 DNA copies in standard material, intact HIV-1 DNA in PLWH was quantified in three different ways (IPDA positive, D4PCR positive and Rainbow positive), resulting in 50 samples with reportable IPDA positive intactness results and 43 samples with reportable D4PCR positive/Rainbow positive intactness results. The median intact copy numbers from the latter 43 samples, quantified by two regions (IPDA), four regions (D4PCR) and five regions (Rainbow) were respectively 26.6, 14.1 and 14.1 copies/million CD4+ T cells before DSI (DNA Shearing Index) correction. As shearing is expected, the DSI (median: 37.3%, IQR: 34.3-38.7%) was used to correct these intact copy numbers, leading to a median of 41.1 (IPDA positive), 22.8 (D4PCR positive) and 22.8 (Rainbow positive) intact HIV-1 copies/million CD4+ T cells. When comparing these intactness results, IPDA intact copy numbers and D4PCR/Rainbow intact copy numbers were significantly different from each other (P-value < 0.0001; Wilcoxon signed-rank test) (*Figure 3e*).

#### Total HIV-1 DNA

Total HIV-1 DNA was detected by the total HIV-1 DNA assay (targeting RU5) in all 69 PLWH on ART. In 64/69 PLWH this resulted in a quantifiable copy number above the LoD95 of 10 copies input, reaching a 92.8% quantification success rate in this cohort. The median total HIV-1 DNA copy number of all samples above the LoD95 (N = 64/69) was 464 copies/million CD4 T cells.

Alternatively, total HIV-1 DNA was defined as the sum of the defective HIV-1 copies (single positives) and intact HIV-1 DNA copies (double positives), in the classical IPDA setup due to the lack of a total HIV-1 DNA readout in previous studies. Here, this resulted in quantifiable results in 50/69 PLWH, as total HIV-1 DNA could only be calculated in samples where the amplification of psi and *env* was successful. In samples with a total HIV-1 DNA (based on RU5) and alternative total HIV-1 DNA based on single and double positives for psi and *env*) read-out (N = 48), the total HIV-1 DNA levels showed a strong correlation (Pearson correlation test, r = 0.97, p < 0.0001) (*Figure 3f*).

#### Proposed decision tree for HIV-1 intactness analysis via Rainbow

Loss of signal in a target region, due to PCR inhibition caused by primer/probe mismatches, can hamper intactness quantification in our multiplex rainbow setup, as is the case for the IPDA setup. To maximize interpretation of intactness results and cope with signal failures we introduce the following strategy (*Figure 4*). Depending on which assay shows a signal failure, we use a decision tree to report on intactness albeit with a lower plexity but keeping the IPDA psi and *env* target regions as a minimal readout for intactness (e.g. 2, 3 or 4 colors). This approach ensures reliable and more precise results without compromising the size of the dataset. By implementing the proposed strategy on this cohort, the dataset increases from 31 to 36 quantifiable intact copy numbers (*Figure 4*). Finally, this 5-color dPCR assay can offer flexibility in the interpretation of different total and/or intactness HIV-1 DNA read-outs if desired (*Table S3*).

**Figure 4:**
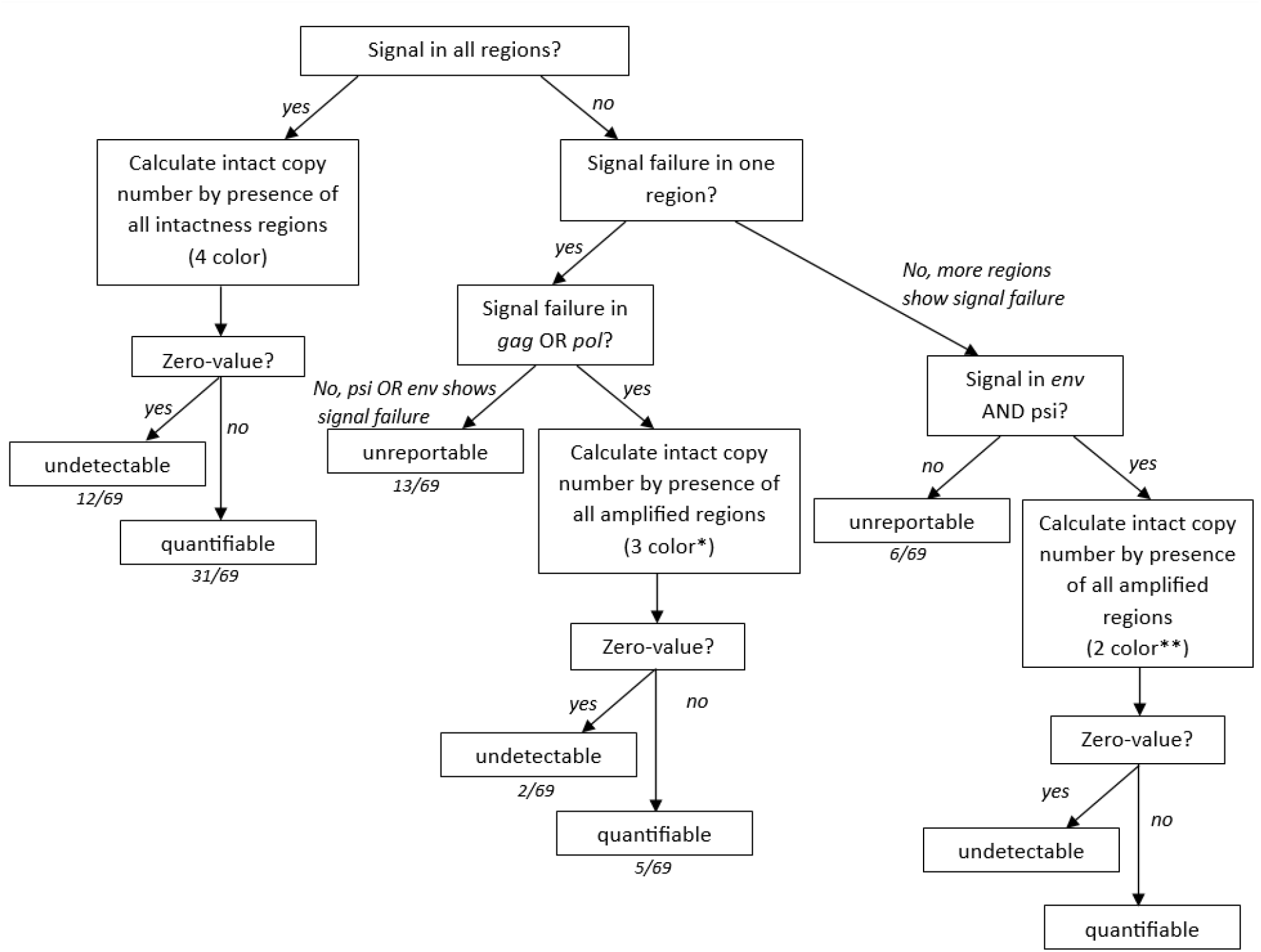
Decision tree for the quantification of intactness levels in PLWH with the Rainbow proviral HIV-1 dPCR assay. If signal failure occurs in certain regions and the intact copy number is calculated by the presence of the amplified regions, it should be noted that this result will be less precise than intact copy numbers calculated by all intactness regions. Intact copy numbers calculated by three regions should be annotated by *. Intact copy numbers calculated by two regions should be annotated by **.

## Discussion

### Rainbow setup and technical performance

To date, new cure strategies are being developed to eliminate the HIV-1 viral reservoir. To evaluate their success and understand HIV-1 reservoir dynamics, accurate and precise measurement of the viral reservoir and particularly its clinically relevant, intact component is essential. Preferentially, this is done with methods that offer sufficient throughput to apply in large(r) clinical trials with high sensitivity and specificity [4, 5, 16].

HIV-1 intactness assays have been published that target specific HIV-1 DNA regions that have the ability to discriminate intact from defective proviruses. These assays were designed based on information on sequence conservation from HIV-1 full-length databases and were heavily cross-validated in previous studies [5, 12]. The IPDA quantifies intact proviruses, using two probes targeting psi and *env* [12]. The Q4PCR approach quantifies intact proviruses by combining qPCR, using four probes, and nFGS analysis [5]. Recent studies showed that Q4PCR underestimates absolute levels of intact proviruses, compared to IPDA due to relatively inefficient PCR amplification of long genomes [16, 17]. Additionally, IPDA misclassifies HIV-1 proviruses that are intact at both psi and *env* but defective elsewhere, leading to an underestimation of efficacy from therapies that target intact proviruses [16].

For this purpose, we developed the Rainbow proviral HIV-1 DNA dPCR assay which is based on a combination of three existing HIV-1 assays to simultaneously measure total HIV-1 DNA and intact HIV-1 proviral DNA with good sensitivity and specificity. This multiplex dPCR approach maximizes information retrieval from the sample (reducing sample input requirements), saves time and offers flexibility as different combinations of PCR assays against HIV-1 regions can be optimized.

The Rainbow proviral HIV-1 DNA dPCR assay was first tested extensively using validated standard material before applying it to samples from PLWH. The results demonstrated that the Rainbow assay is sensitive up to 10 copies HIV-1 input per reaction, shows good linearity and is reproducible in all five multiplexed assays. In addition, clear separation is observed of the positive partitions in all channels and combinations, indicating PCR compatibility of the assays. Fluorescence spill-over between dPCR channels in this assay was limited to the red channel into the orange channel, without hampering the separation of the positive partitions. As expected, the variation (CV%) increased in the lower input range but is well within acceptable levels [21]. Finally, there is limited overquantification as higher HIV-1 DNA levels were observed than theoretically estimated across the standard curve. This can be explained by the fact that calculations of input material were made based on DNA concentration measured by fluorometric quantification (Qubit) and the assumption that the mass of one tetraploid J-Lat cell is 13.2 *p*g. These results are in line with a recent study where the classical IPDA was adapted for the QIAcuity 5-plex system [22].

### HIV-1 intactness in PLWH: IPDA performance

Since the IPDA was published, several follow-up studies have reported on failure rates of 6.3-28% due to underlying sequence diversity/polymorphisms [23–27]. If we assess the IPDA setup in our study, signal failure in both psi and *env* regions was observed in 4.3% (3/69) of the samples and in one of the two regions in 23.2% (16/69) of the samples, corroborating previous reports on IPDA performance in HIV-1 subtype B.

### HIV-1 intactness in PLWH: D4PCR performance

Including additional regions in our Rainbow setup, allowed for a better and more robust estimate of intactness levels as proviruses are excluded that are intact in the psi and *env* region, but defective in the *pol* and/or *gag* region. When comparing intactness estimates with 4 regions, we found significantly lower values in 31/46 PLWH than when only considering two regions (IPDA), indicating the added value.

In this cohort, a loss of signal was observed in one or more primer/probe sets from the Rainbow proviral HIV-1 DNA dPCR assay in 26/69 (37.7%) of the samples which is in line with a previous study examining the Q4PCR intactness assay. In that study, 6/9 patients showed HIV-1 polymorphisms that caused a loss of signal for at least one of the primer/probe sets from the Q4PCR assay [5].

Overall, it remains difficult for intactness assays when signal failures are observed, to fully address the nature of this phenomenon. For this, parallel full-length sequencing could reveal potential causes of PCR failure such as primer/probe mismatches. In addition, when using 2-region intactness readouts, failure of one assay inhibits estimation of intact HIV-1 DNA. This can be countered by increasing the number of regions, as described by the Rainbow proviral HIV-1 DNA assay, Q4PCR, 5T-IPDA and CS-IPDA. Depending on which primer/probe set that resulted in signal failure, intactness levels can still be calculated by the remaining amplified regions as proposed in our decision tree. For example, an earlier study reported signal failure in the *env* region for a PLWH, but several intact proviral sequences scored positive for psi, *pol* and *gag* by Q4PCR and were confirmed by nFGS sequencing [5]. However, this still can lead to an overestimation of the intact viral reservoir size.

### Total HIV-1 DNA in PLWH

Multiple assays for total HIV-1 DNA quantification have been described previously. We selected the RU5 assay [18, 19], as it was one of the top performing total HIV-1 DNA assays, according to the benchmarking study of Rutsaert et al., that captures on average 93% of the samples by ddPCR [11]. In our cohort, the RU5 assay performed equally well in the 5-plex Rainbow dPCR setup with 92.8% of the samples having a quantifiable total HIV-1 DNA level above the LoD95 of 10 copies per reaction.

In some studies, using the classical IPDA 2-plex setup, total HIV-1 DNA is quantified as the sum of the intact and defective proviruses without DSI correction [26, 28]. However, these regions are prone to a higher failure rate, as compared with total HIV-1 DNA assays. In our study, the total HIV-1 DNA results, calculated by the sum of intact and defective proviruses (IPDA setup), could be obtained in 72.5% of the samples, whereas the total HIV-1 DNA assay was successful in 92.8% of the samples. This demonstrates the added value of including a dedicated total HIV-1 DNA assay in the Rainbow proviral HIV-1 DNA dPCR assay.

### Limitations

The Rainbow assay targets five relatively small regions from the ∼9 kb HIV-1 genome. Therefore, the Rainbow assay will likely still overestimate the true viral intact reservoir size, as all PCR-based approaches. In terms of dPCR data analysis, patient specific thresholding was mandatory, increasing the analysis time. In addition, setting up 5-plex assays, proper spill-over correction is needed, as it further challenges the thresholding procedure due to tilting of partition populations.

As described before, unreportable results were observed in 37.7% of the PLWH in this cohort, probably due to mismatches in the target region. This is higher than the 27.5% failure rate by IPDA, resulting in a smaller but more accurate dataset for intactness analysis. With the proposed strategy to quantify intactness levels with at least two intactness regions and up to four regions, a balance is achieved between the size of the dataset and the precision of intactness levels. Moreover, different readouts for intactness are possible with the Rainbow proviral HIV-1 DNA assay.

In case of a low viral reservoir size, it is possible that no HIV-1 DNA is detected because too low gDNA input was used. This can be obviated by increasing the input but does not guarantee success. Therefore, interpretation of intactness results remains difficult and supporting full-length sequencing data can help to address primer/probe failures. Additionally, the performance of the Rainbow assay was only assessed in subtype B PLWH. Therefore, a more extensive study should be performed to confirm that the target regions from the Rainbow DNA assay can quantify intact HIV-1 DNA in other subtypes.

Similar to quantification results by the classical IPDA, intact copy numbers quantified by the Rainbow assay, need to be corrected for DNA shearing. As this might inflate real intactness quantification, we advocate to report intactness levels with and without DNA shearing correction. In addition, the formula published by Bruner et al. used for DNA shearing correction requires an additional dilution to assess DNA shearing on the RPP30 gene and does not take into account the distribution of the single positive partitions for each HIV-1 assay that are present in the sample. Future research should try and exclude this mandatory dilution and establish a DNA shearing correction formula that takes into account the abundance of single positive partitions for HIV-1 specific regions.

This Rainbow approach shows the flexibility of combining PCR assays for different quantification purposes and maximize the retrieval of information on the reservoir while reducing cost and amount of biological material. This rainbow approach is not restricted to these 5 regions and can be modified to the researcher’s interest. With future developments of dPCR platforms or higher order multiplexing strategies, even more regions could be combined to reach an intactness quantification.

In conclusion, the Rainbow proviral HIV-1 DNA dPCR assay shows the necessary throughput to be implemented in larger clinical trials and renders information on total HIV-1 DNA levels and intact HIV-1 DNA levels. By combining four intactness regions, comparable sensitivity and enhanced specificity is obtained (compared to IPDA) without introducing bias against full-length proviruses. Overall, it gives a better estimate of the true viral reservoir size which is of benefit to all clinical trial studies.

## Material and methods

### Ethics statement

Study participants provided written consent for this study. The participants are all HIV-1 infected individuals on suppressive ART with an undetectable viral load of < 50 copies/mL HIV-1 RNA for at least 3 months (*NCT04553081)*.

### Sample preparation

#### J-Lat type 10.6 dilution series

J-Lat cells 10.6 (containing one HIV-1 subtype B copy per cell) were cultured in complete RPMI (37°C, 5% CO2), containing Glutamax, PenStrep and Foetal calf serum (FCS). Dry cell pellets were generated by centrifugation (500g, RT) and stored at −80°C.

Peripheral blood mononuclear cells (PBMCs) from 8 HIV-1 negative individuals were pooled after isolation by densitiy gradient centrifugation. CD4+ T cells were isolated by negative selection using the EasySep CD4 T cell isolation kit (Stemcell Technologies) and frozen as dry pellet at −80°C.

For the preparation of the 12-step two-fold limiting dilution curve, genomic DNA (gDNA) was extracted from J-Lat cells 10.6 and HIV-1 negative CD4+ T cells with the QIAcube (Qiagen), using the QiaAmp DNA mini kit (Qiagen). CD4+ T cell gDNA was used to establish negative template controls and as a background matrix to dilute J-Lat gDNA to an estimated 512, 256, 128, 64, 32, 16, 8, 4, 2, 1, 0.5 and 0.25 copies/µl. The two-fold dilution curve was prepared at three different timepoints by the same person. Aliquots were stored at −20°C.

#### Samples from PLWH

CD4+ T cells were isolated by negative selection, starting from freshly isolated PBMCs from 69 PLWH using the EasySep CD4 T cell isolation kit (Stemcell Technologies). CD4+ T cells were frozen as dry pellet at −80°C prior to DNA extraction with the DNeasy Blood & Tissue kit (Qiagen). Extracted gDNA was stored at −20°C.

### Rainbow proviral HIV-1 DNA dPCR assay

The Rainbow proviral HIV-1 DNA dPCR assay is a multiplex dPCR approach that combines five different HIV-1 regions to quantify total HIV-1 DNA and intact HIV-1 DNA simultaneously. Primers and probe sequences were acquired from Bruner et al. (2019) [12], Gaebler et al. (2019) [5], Yun et al. (2002) [18] and Yu et al. (2008) [19] *(Figure S1, Table S1).* For the set-up of the Rainbow proviral HIV-1 DNA dPCR assay, the same strategy was used as van Snippenberg et al. (2021) [29]. The *gag* probe was modified to contain a Cy5 fluorophore while maintaining the psi probe (linked to FAM fluorophore) and the *env* probe (linked to HEX fluorophore).The dark competition probe for the *env* region was modified with LNAs to increase the melting temperature instead of adding a minor groove binder (MGB). The *pol* probe and total (RU5) probe were modified to contain the ATTO550 fluorophore and the ROX fluorophore, respectively. All sequences were ordered and synthesized by Integrated DNA Technologies (IDT).

In general, the Rainbow proviral HIV-1 DNA dPCR assay was performed on a QIAcuity Four digital PCR platform (Qiagen, Germany) with at least 500 ng (CD4+ T cell) gDNA input per reaction. The reaction mix preparation for the Rainbow proviral HIV-1 DNA assay was as follows: 10 µL 4x concentrated, ready to use Qiacuity Probe Master Mix (ID: 250102, Qiagen), 2 µL of each primer/probe set (*Table S1*), 0.3 µL restriction enzyme XBaI (100,000 units/ml), 4 µL template gDNA and nuclease free water to 40 µL total volume per reaction. The PCR reaction mixes were transferred into a 26K 24-well nanoplate (ID: 250001, Qiagen). Partitioning of the PCR mix was performed on the QIAcuity Four digital PCR system after sealing the plate. The digital PCR program started with 2 minutes at 95°C for the activation of the Qiacuity probe mix, followed by 40 cycles of 94°C for 30 seconds and 56°C for 60 seconds. Imaging settings were 500 ms, 500 ms, 400 ms, 200 ms and 400 ms in the green, yellow, orange, red and crimson channel, respectively.

The three standard dilution curves were measured in 20 replicates for each dilution point and a total of 288 NTCs were measured to establish the limit of blank (LoB). Patient samples were run in triplicate with on each plate two high positive controls, two low positive controls and two negative controls. These control samples were prepared similarly to the dilution curve.

### RPP30 assay

The RPP30 duplex assay was performed as described by Bruner et al. and used to normalize the total and intact HIV-1 DNA copy numbers to the cell number input. In addition, the DNA shearing index (DSI) was calculated and used to correct the intact HIV-1 DNA copy number. For this assay, gDNA was diluted 100 times before adding it to the reaction mix, as each CD4+ T cell contains two copies of the RPP30 gene. Primers and probes were used as described by Bruner et al. (2019) [12]. The same digital PCR thermal cycling program was used as for the Rainbow proviral HIV-1 DNA dPCR assay. Samples were run in duplicate with two positive controls (J-Lat gDNA spiked into HIV-1 negative CD4+ T cell gDNA) and two negative controls (nuclease free water).

### Data analysis

Raw data was analyzed with QIAcuity Software Suite version 1.2. Thresholding was done manually based on positive and negative controls, while taking into account spill-over from adjacent channels (red into orange). For the standard material, the same manual threshold was used per plate, whereas for patient samples the manual thresholding was adapted if needed as these cross-plate thresholds did not always fit individual samples. Intact HIV-1 DNA was quantified based on two regions from the classical IPDA (psi, *env*) and based on four (D4PCR) or five regions from the Rainbow proviral HIV-1 DNA dPCR assay if possible. Finally, intact HIV-1 DNA levels were adjusted for DNA shearing and normalized per million CD4+ T cells via the RPP30 duplex assay as previously described [12].

### Statistical analysis

Statistical analysis and figures were made in Rstudio (2023.06.01 Build 524 “Mountain Hydrangea” release) running R (v. 4.3.0), GraphPad Prism 9.5.1 and Biorender (www.Biorender.com). The R shiny tool dPCalibRate (v1.1) was used to evaluate the results of the performance of the Rainbow assay on the standard curve [30].The non-parametric Wilcoxon signed-rank test was performed to assess whether the average difference of intactness levels defined by two regions and four regions was significantly different from 0. Pearson correlation coefficient was calculated to test the correlation between the total HIV-1 DNA levels obtained by the total HIV-1 DNA assay and by the sum of the IPDA intact and defective proviruses. Finally, Spearman correlation coefficients were used to check the associations between virological and clinical parameters.

## Acknowledgments

We thank all study participants who donated blood samples, as well as MDs and study nurses who helped with the recruitment and coordination of this study.

## Funding

- Fund for scientific research (FWO JUNIOR: G0B3820N)
- SBO-SAPHIR (S000319N)
- ViiV Healthcare (A21/TT/1555)

### Author contributions

Conceptualization: MD, WVS, WT
Methodology: MD, WVS, WT
Investigation: MD, WVS, MV, EDS
Formal analysis: MD, WT
Supervision: WT, EEB, SR, SG, LV
Resources: MADS, LV
Funding acquisition: LV
Writing—original draft: MD
Writing—review & editing: MD, WVS, EEB, SR, SG, LV, WT

### Data and materials availability

All data needed to evaluate the conclusions in the paper are present in the paper and/or the Supplementary Materials.

