## Supplemental material for "Integrative assessment of total and intact HIV-1 reservoir by a five-region multiplexed Rainbow digital PCR assay"

Mareva Delporte *et al.*

| RAINBOW | Sequence | HXB2 | Conc (nM) | Reference |
| --- | --- | --- | --- | --- |
| RU5 Fwd | TTAAGCCTCAATAAAGCTTGCC | 518 🡪 539 | 675 | Yun et al (2002) |
| RU5 Rvd | GTTCGGGCGCCACTGCTAGA | 628 🡪 647 | 675 |  |
| RU5 probe | /56ROXN/CCAGAGTCACACAACAGACGGGCACA/3IAbRQSp/ | 559 🡨 584 | 187.5 | Yu et al (2008) |
| psi Fwd | CAGGACTCGGCTTGCTGAAG | 692 🡪 711 | 675 | Bruner et al (2019) |
| psi Rvd | GCACCCATCTCTCTCCTTCTAGC | 775 🡨 797 | 675 |  |
| psi probe | /56-FAM/TTTTGGCGT/ZEN/ACTCACCAGT/3IABkFQ/ | 740 🡨 758 | 187.5 |  |
| *env* Fwd | AGTGGTGCAGAGAGAAAAAAGAGC | 7736 🡪 7759 | 500 |  |
| *env* Rvd | GTCTGGCCTGTACCGTCAGC | 7832 🡨 7851 | 500 |  |
| *env* dark probe | CC+TTAGGTTCTTAGG+AGC | 7781 🡪 7798 | 250 |  |
| *env* probe | /5HEX/CCTTGGGTT/ZEN/CTTGGGA/3IABkFQ/ | 7781 🡪7796 | 250 |  |
| *gag* Fwd | ATGTTTTCAGCATTATCAGAAGGA | 1300 🡪 1323 | 337.5 | Gaebler et al (2019) |
| *gag* Rvd | TGCTTGATGTCCCCCCACT | 1359 🡨 1377 | 337.5 |  |
| *gag* probe | /5Cy5/CCACCCCAC/TAO/AAGATTTAAACACCATGCTAA/3IAbRQSp/ | 1325 🡪 1354 | 93.75 |  |
| *pol* Fwd | GCACTTTAAATTTTCCCATTAGTCCTA | 2536 🡪 2562 | 675 |  |
| *pol* Rvd | CAAATTTCTACTAATGCTTTTATTTTTTC | 2634 🡨 2662 | 675 |  |
| *pol* probe | /5ATTO550N/AAGCCAGGAATGGATGGCC/3IAbRQSp/ | 2586 🡪 2604 | 187.5 |  |

**Table S1**: Primer and probe sequences of each target region in the Rainbow proviral HIV-1 DNA dPCR assay with their respective location in the HXB2 genome, the concentration used and the corresponding publication for each target region.

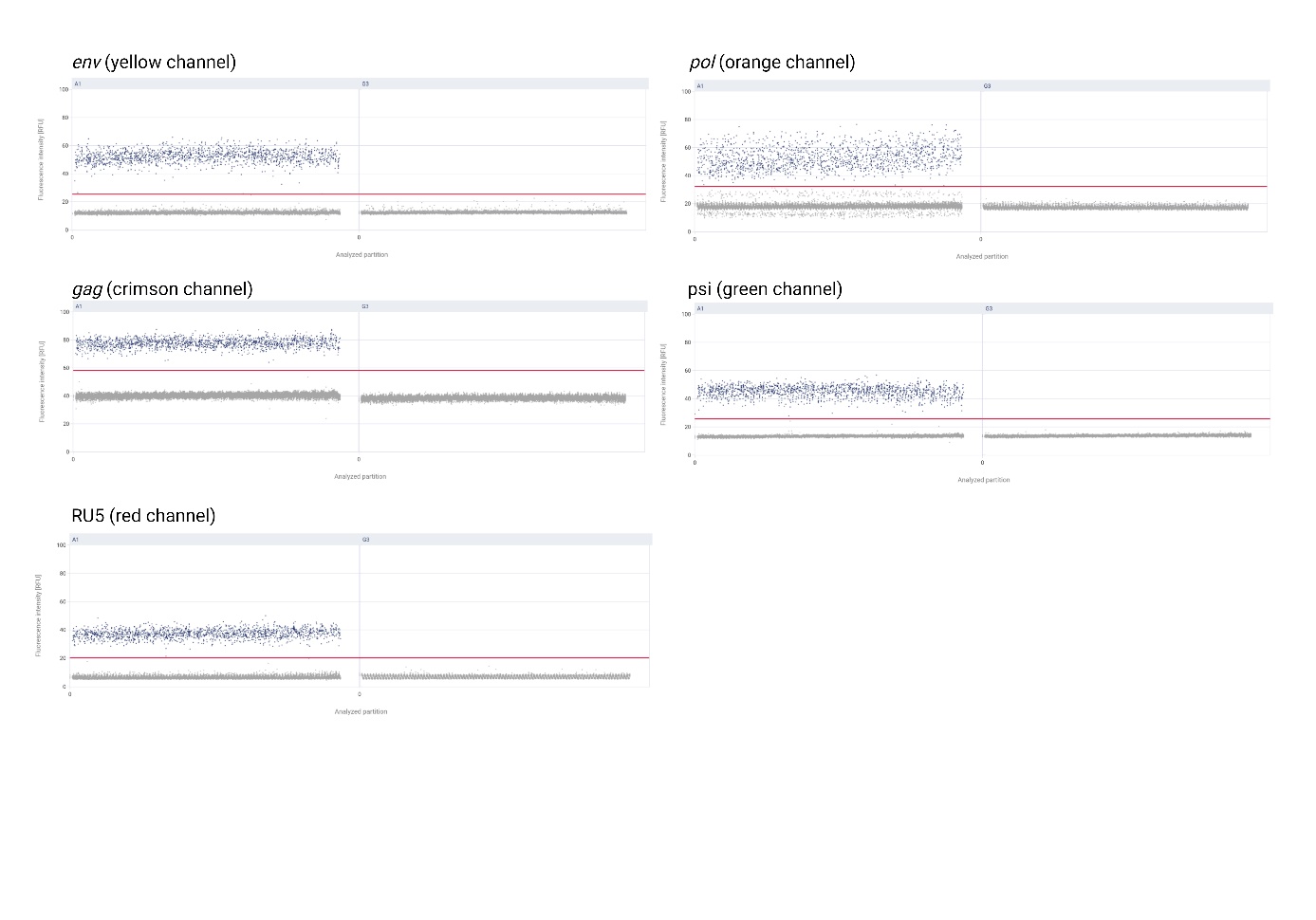

**Figure S1:** 1D-scatterplots for each target region in QIAcuity Software Suite 2.1.7.182 with raw data of HIV-1 copies (J-Lat gDNA) spiked into HIV-1 negative CD4 background gDNA (A1) and raw data of HIV-1 negative CD4 background gDNA (G3). Red line represents the threshold for each target region.

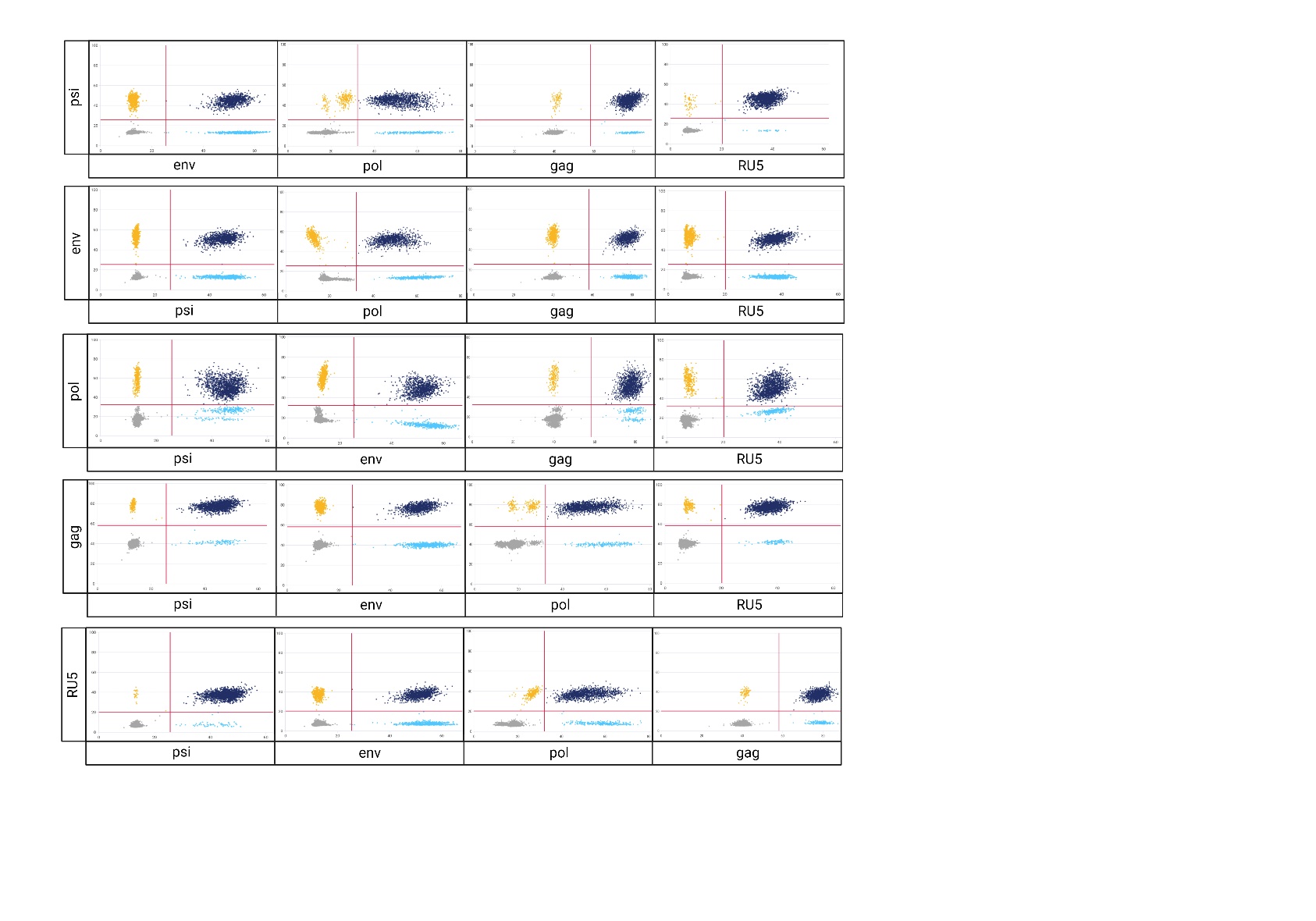

**Figure S2**: 2D-scatterplots for all dual combinations of target regions in the QIAcuity Software Suite 2.1.7.182 with raw data of HIV-1 copies (J-Lat gDNA) spiked into HIV-1 negative CD4 background gDNA. In each 2D-scatterplots, the fluorescence intensity (RFU) of a target region is plotted against the fluorescence intensity (RFU) of another target region. The red lines represent the threshold for each target region.

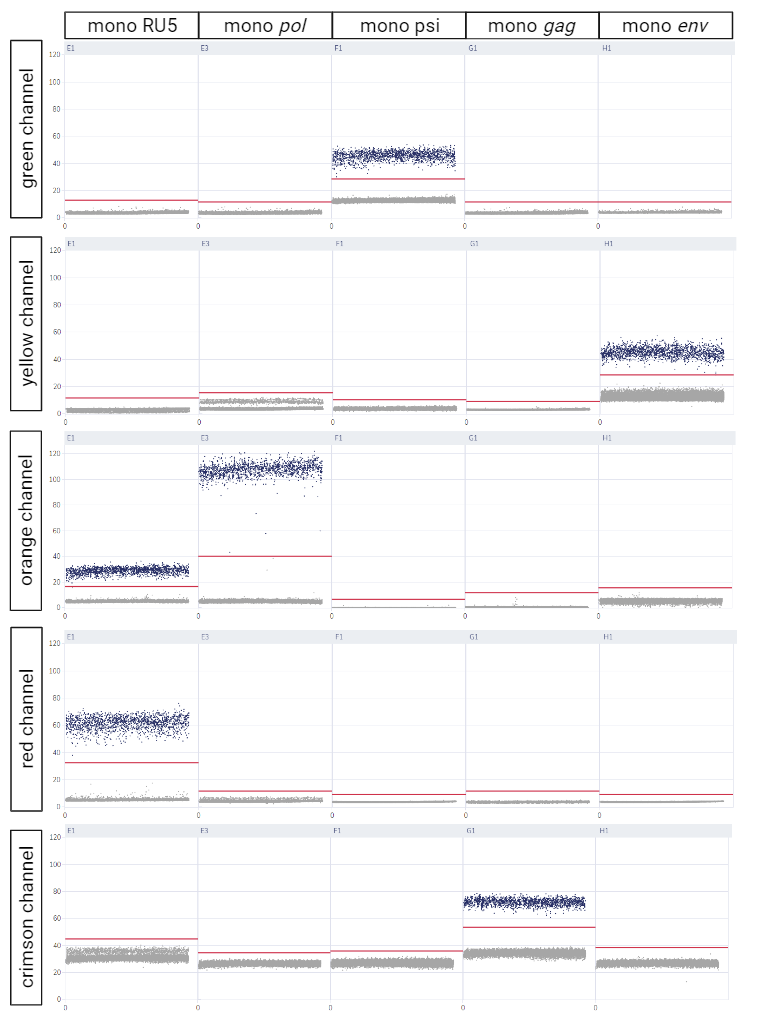

**Figure S3:** 1D-scatterplots of monocolor assays generated in the Qiacuity Software Suite. Each column represents the 1D scatterplots of the monocolor assays in each fluorescence channel (row).The red lines represent the thresholds for each target region. In the yellow fluorescent channel, limited spill-over is observed from the orange fluorescent channel. In the orange and crimson fluorescent channel, spill-over is observed from the red fluorescent channel.

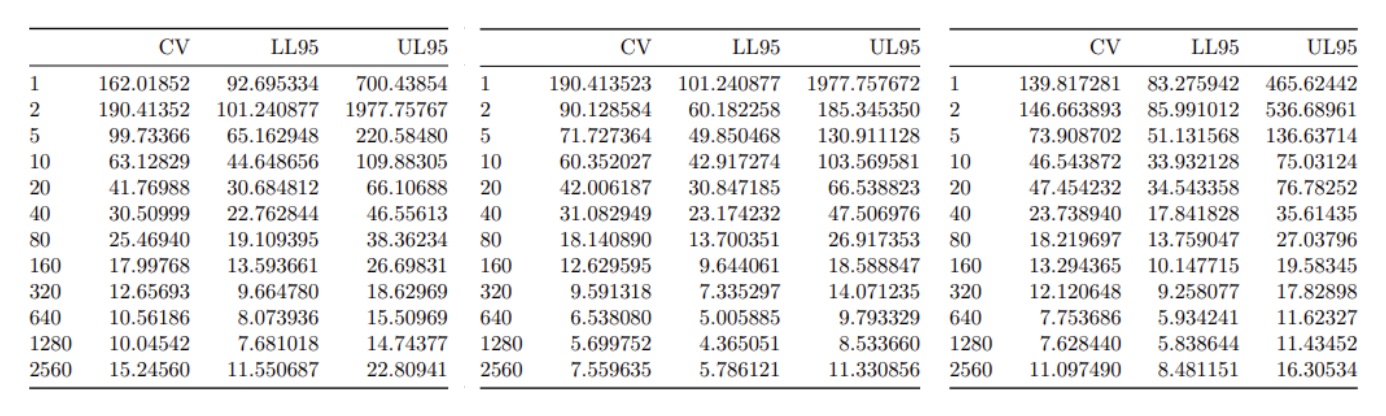
**Table S2:** Coefficiënts of variance for each dilution point per standard curve generated by dPCalibRate v1.1

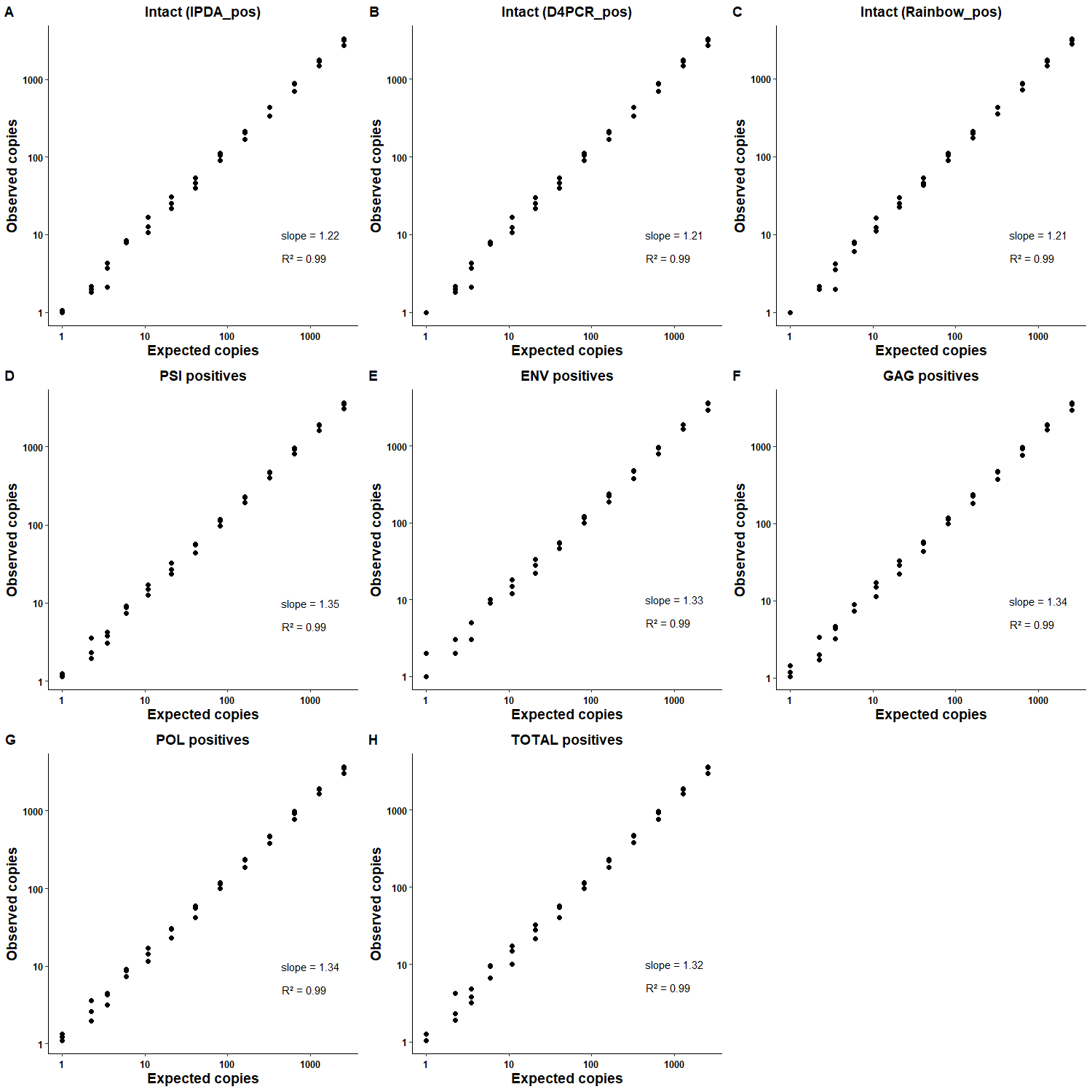

**Figure S4: Linearity and accuracy of Rainbow proviral HIV-1 DNA dPCR assay on HIV-1 standard material (J-Lat gDNA spiked into CD4+ gDNA).**

Three standard dilution curves were prepared to achieve predicted inputs of 0 to 2560 copies (x axis). The observed copies of IPDA psi+*env*+ (A), psi+*env*+*gag*+*pol*+ (B), psi+*env*+*gag*+*pol*+RU5+ (C) corrected for shearing; psi+ (D), *env*+ (E), *gag*+ (F), *pol*+ (G) and RU5 (total)+ (H) were measured by the Rainbow proviral HIV-1 DNA dPCR assay (y-axis). Each dot represents 20 replicates of one standard curve. Slopes and R² were determined by linear regression analysis.

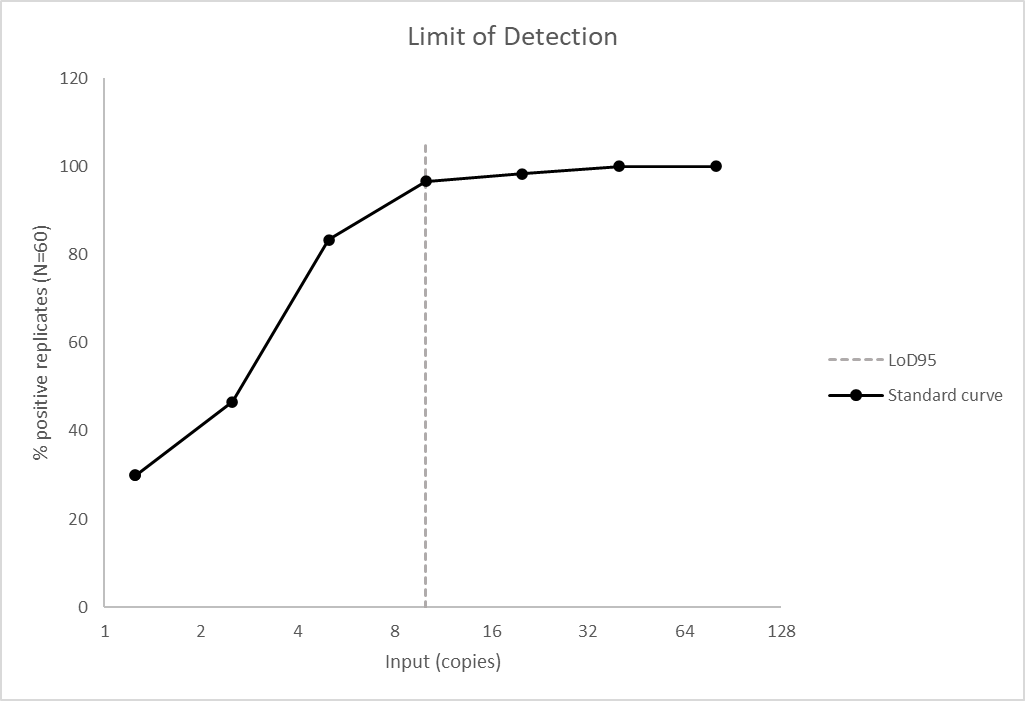

**Figure S5:** **Limit of detection curve for all three standard curves in lowest dilution points (range 1.25-80 copies input).**

The positivity rate of all replicates (n=60) is plotted for each dilution point, ranging from 1.25 copies input to 80 copies input. This curve applies to the positivity rate of psi, *env*, *gag*, *pol*, RU5, psi+*env*+, psi+*env*+*gag*+*pol*+ and psi+*env*+*gag*+*pol*+RU5+.

**Table S3: Overview of different possible read-outs with the Rainbow proviral HIV-1 DNA assay.** Total HIV-1 DNA can be measured by the dedicated total HIV-1 DNA assay (RU5) or by the presence of the other target regions. Intact HIV-1 DNA can be quantified by 2, 3, 4 or 5 target regions.

| **Total HIV-1 DNA** | **Intact HIV-1 DNA** |
| --- | --- |
| RU5 |  |
| psi, *env* | IPDA (psi AND *env*) |
| psi, *env*, *pol* | TRIPGAG (psi AND *env* AND *gag*) |
| psi, *env*, *gag* | TRIPPOL (psi AND *env* AND *pol*) |
| psi, *env*, *gag*, *pol* | D4PCR (psi AND *env* AND *gag* AND *pol*) |
| psi, *env*, *gag*, *pol*, RU5 | RAINBOW (psi AND *env* AND *gag* AND *pol* AND RU5) |
